# A *Drosophila* model for Costello Syndrome caused by Ras mutation K117R

**DOI:** 10.1101/2025.02.03.635762

**Authors:** Prashath Karunaraj, Emily Cao, Harnoor Singh, Max Luf, Stacey J. Baker, E Premkumar Reddy, Cathie M. Pfleger

**Affiliations:** Department of Oncological Sciences The Icahn School of Medicine at Mount Sinai New York, NY 10029; The Tisch Cancer Institute The Icahn School of Medicine at Mount Sinai New York, NY 10029; The Graduate School of Biomedical Sciences The Icahn School of Medicine at Mount Sinai New York, NY 10029; Department of Medicine Columbia University, Irving Medical Scenter New York, NY 10032

**Keywords:** Ras, *Drosophila*, Costello Syndrome, RASopathies, Trametinib, Rigosertib

## Abstract

Germline mutations that increase signaling through the Ras pathway can cause developmental disorders called RASopathies. The RASopathy Costello syndrome has been described to present with hallmarks that include short stature, intellectual disability, cardiac issues, and characteristic facial abnormalities and has been associated with gain-of-function mutations in HRas. The most common HRas mutations in Costello Syndrome occur at G12 and G13, but there are also other rare mutation sites such as K117 including HRas^K117R^. Ras^K117R^ mutations are also found in colorectal cancer. *Drosophila* studies modeling gain-of-function in Ras primarily utilize the common cancer-associated mutation G12V, and previous *Drosophila* RASopathy models assessing Ras gain-of-function mutations have used human sequences for KRas G12D and HRas G12S. To augment these studies, we characterized the phenotype of engineering the rare gain-of-function mutation K117R in the *Drosophila* Ras sequence. We report here that constitutive low-level expression of Ras^K117R^ increased lethality and reduced body size while also causing rough eye and ectopic wing vein phenotypes in those flies that survived to adulthood. Ras pathway inhibitors Trametinib and Rigosertib suppressed the lethality but not the reduced size phenotypes. Trametinib strongly suppressed the K117R wing vein phenotype whereas Rigosertib had only subtle effects. Trametinib is a direct MEK inhibitor. Rigosertib has been reported to have strong effects on PI3K signaling and to indirectly inhibit the Raf-ERK branch. Therefore, this data is consistent with an interpretation that some lethality in the fly Ras^K117R^ model depends on elevated signaling through the Raf-ERK branch and potentially some lethality depends on the PI3K branch. In contrast, the lack of effects on the reduced size phenotypes would be consistent with small stature resulting from Raf- and PI3K-independent processes. We propose that this model can be useful for future mechanistic analysis and pharmacological screening and evaluation.

## INTRODUCTION

The important signaling molecule Ras is involved in regulating proliferation, cell survival, cell differentiation, and patterning. Therefore, restricting Ras activation within a healthy range is crucial to normal development and patterning. Ras normally becomes activated by upstream receptor tyrosine kinases (RTKs) such as EGFR after they become activated by binding their ligands. Upon activation by these upstream RTKs, Ras undergoes exchange of GDP to become Ras-GTP and switches to an active conformation. Excess activation of Ras in development can disrupt patterning and lead to a range of consequences. Such excess Ras signaling can occur through a number of mechanisms such as amplification or mutational activation of upstream activators like the RTKS, loss-of-function mutations in inhibitors of Ras signaling such as Neurofibromin (NF1), amplification or gain-of-function mutations in Ras itself, or activating mutations in downstream effectors such as Raf. Such genetic lesions can occur in differentiated cells in adult organisms which can lead to tumorigenesis and therefore have been associated with a range of cancers. In contrast, germline mutations affect every cell in a developing organism and have been associated with a variety of syndromes collectively called RASopathies that include Costello Syndrome, Neurofibromatosis Type 1 (NF1), Noonan’s syndrome, Leopard’s syndrome, and others [Der Kaloustian et al., 1991; Martin et all., 1991; Dard et al., 2022; Kerr et al., 2006. Gelb and Targaglia, 2006; Gelb et al., 2022; Tidyman et al., 2009; Gripp et al., 2012].

*Drosophila* models have been developed to model several RASopathies including by modulating *neurofibromin* (*Nf1*) [The et al., 1997] to model NF1, *corkscrew* (*csw*) to model Noonan’s syndrome [Oishi et al., 2006], Dsor1/MEK to model cardio-facio cutaneous syndrome [Goyal et al, 2017], and transgenic expression of disease associated mutations in human Ras and Raf sequences [Das et al., 2021] among others. Specifically, previous *Drosophila* RASopathy models assessing gain-of-function mutations in Ras itself have utilized human sequences of KRas G12D and HRas G12S [Das et al., 2021].

In *Drosophila*, one Ras gene (referred to as *Ras85D*, *Ras1*, and most commonly referred to simply as Ras) here referred to as Ras corresponds to mammalian Ras isoforms K-Ras, H-Ras, and N-Ras. *Drosophila* are an excellent model in which to model the phenotypes of specific RASopathy-associated variants without complicating the interpretation conferred by isoform-specific differences. Here we report a new Costello syndrome model using an inducible *Drosophila* Ras^K117R^ transgene. Constitutively expressing Ras^K117R^ increased lethality, reduced organism size, and caused rough eyes and ectopic wing veins. We show that this model can be used to evaluate the efficacy of pharmacological intervention. MEK inhibitor Trametinib increased survival to adulthood and suppressed the ectopic wing vein phenotypes but did not suppress the reduced organ size phenotypes; in fact, Trametinib enhanced the small, rough eye phenotype. Rigosertib also increased survival to adulthood but did not suppress the organ size phenotypes.

## RESULTS AND DISCUSSION

### Constitutive Ras^K117R^ expression increased lethality and reduced organism and organ size

We created a UAS transgene of FLAG-His6 tagged Ras^K117R^ (see methods for additional details) which can be expressed using the Gal4 system. To model the constitutive expression throughout all cells in a developing animal as would be directed by germline mutations, we utilized constitutive low-level gal4 driver *Act5C-gal4*. Expressing Ras^WT^ [Washington et al., 2020] with *Act5C-gal4* (*Act5C>Ras^WT^*) resulted in survival rates similar to *Act5c-gal4* controls (*Act5C-gal4/+*) (Fig. 1A). In contrast, expressing Ras^K117R^ with *Act5C-gal4* (*Act5C>Ras^K117R^*) resulted in a dramatic decrease in pupal survival to adulthood (Fig. 1A). Lethality appears to be increased in males; in several trials, flies that survived to adulthood were primarily female although this was not always statistically significant due to the low number of surviving flies overall (Fig. 1A’).

**Figure 1:**
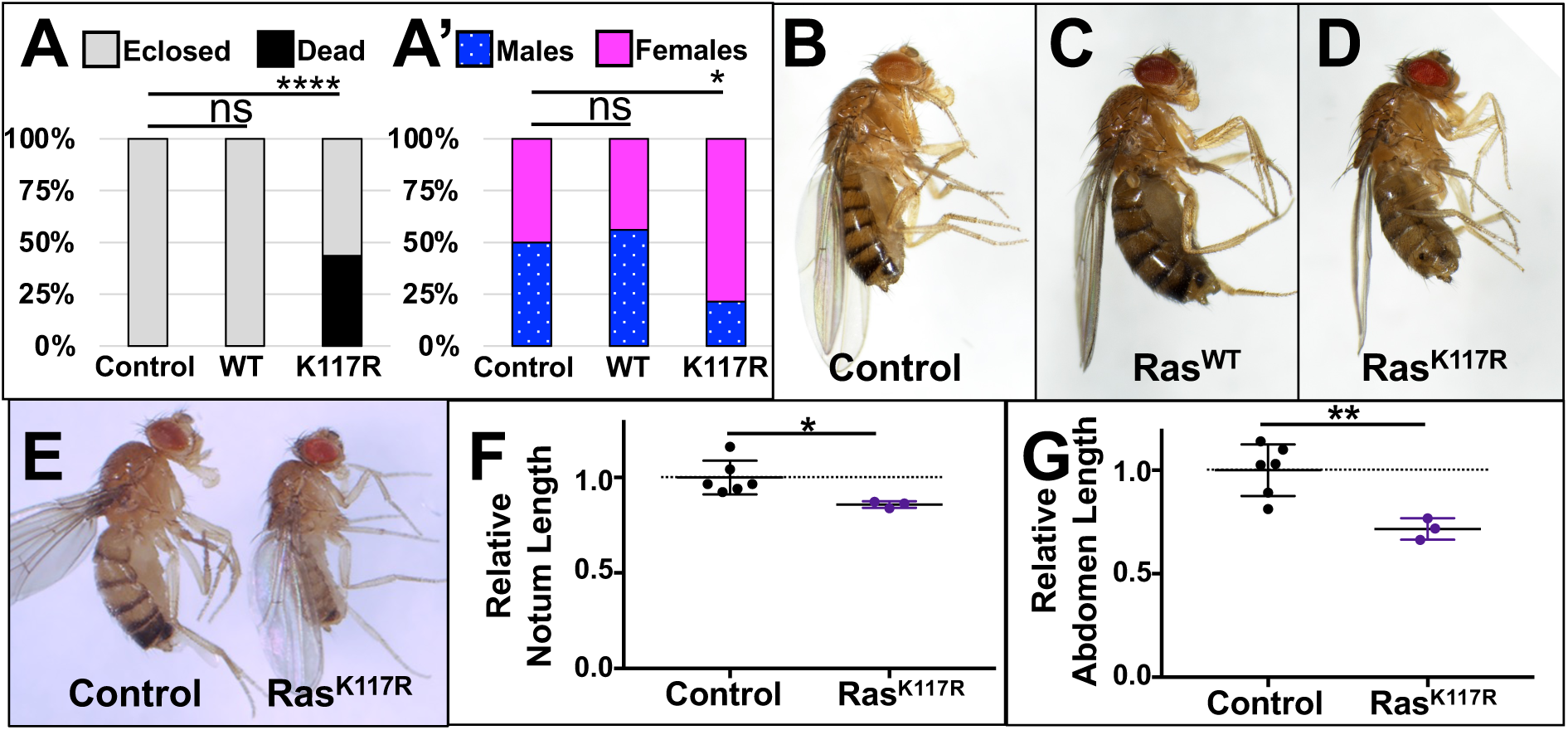
Constitutive expression of Ras^K117R^ reduces survival and organism size. (A) Bar graph summarizing relative proportions of pupae to survive (gray/upper portion of each bar) or die (black, lower portion of each bar) at 21°C. *Act5C-gal4/+* control pupae (lane 1) generally survived to adulthood with little to no lethality. *Act5C>Ras^WT^* pupae (lane 2) showed no statistically different survival compared to control pupae. *Act5C>Ras^K117R^*pupae (lane 3) showed a dramatic decrease in survival to adulthood. Reproducibly, only 25-40% of *Act5C>Ras^K117R^* pupae eclose (example of a trial with lower survival shown later in Fig. 3A). (A’) Bar graph summarizing the relative proportions of adult males and females of the relevant genotypes that eclose. Males (spotted blue, lower portion of bars) were relatively equal to the number of females (solid pink, upper portion of bars) that reached adulthood for *Act5C-gal4/+* control flies (lane 1) and *Act5C>Ras^WT^* flies (lane 2). In contrast, far fewer Act5C>Ras^K117R^ males survive relative to the number of females, although this was not always statistically significant due to the low number of surviving *Act5C>Ras^K117R^*flies. (B) *Act5C-gal4/+* control fly. (C) *Act5C>Ras^WT^*fly. (D) *Act5C>Ras^K117R^* fly. (E) Side-by-side photo of *Act5C-gal4/+* and *Act5C>Ras^K117R^* flies highlighting the size difference. (F) Scatter plot showing the relative notum length of control *Act5C-gal4/+* flies (left) and *Act5C>Ras^K117R^* flies (right). (G) Scatter plot showing the relative abdomen length of control *Act5C-gal4/+* flies (left) and *Act5C>Ras^K117R^*flies (right). Females are shown in panels B-E and analyzed in panels F-G. Dotted lines overlaid on graphs in in F-G indicate the mean for the control. For graphs in A, A’, F, and G ns indicates not significant, *, **, and **** indicate statistically significant in Chi square tests (A, A’) or t tests (F, G); for p values, see Supplemental File S1.

Constitutive Ras^WT^-expressing flies were not obviously different from *Act5C-gal4/+* control flies in terms of gross morphology and overall size (Fig. 1C compared to Fig. 1D), eye area (Fig. 2A), and wing area (Fig. 2B). In contrast, Ras^K117R^-expressing flies that survived to adulthood showed obvious reduction in overall body size (Fig. 1D compared to Fig. 1B, side-by-side example in Fig. 1E), notum length (Fig. 1F), abdomen length (Fig. 1G), eye size (Fig. 2A), and wing size (Fig. 2B). Reduced body size is consistent with the small stature associated with Costello Syndrome.

**Figure 2:**
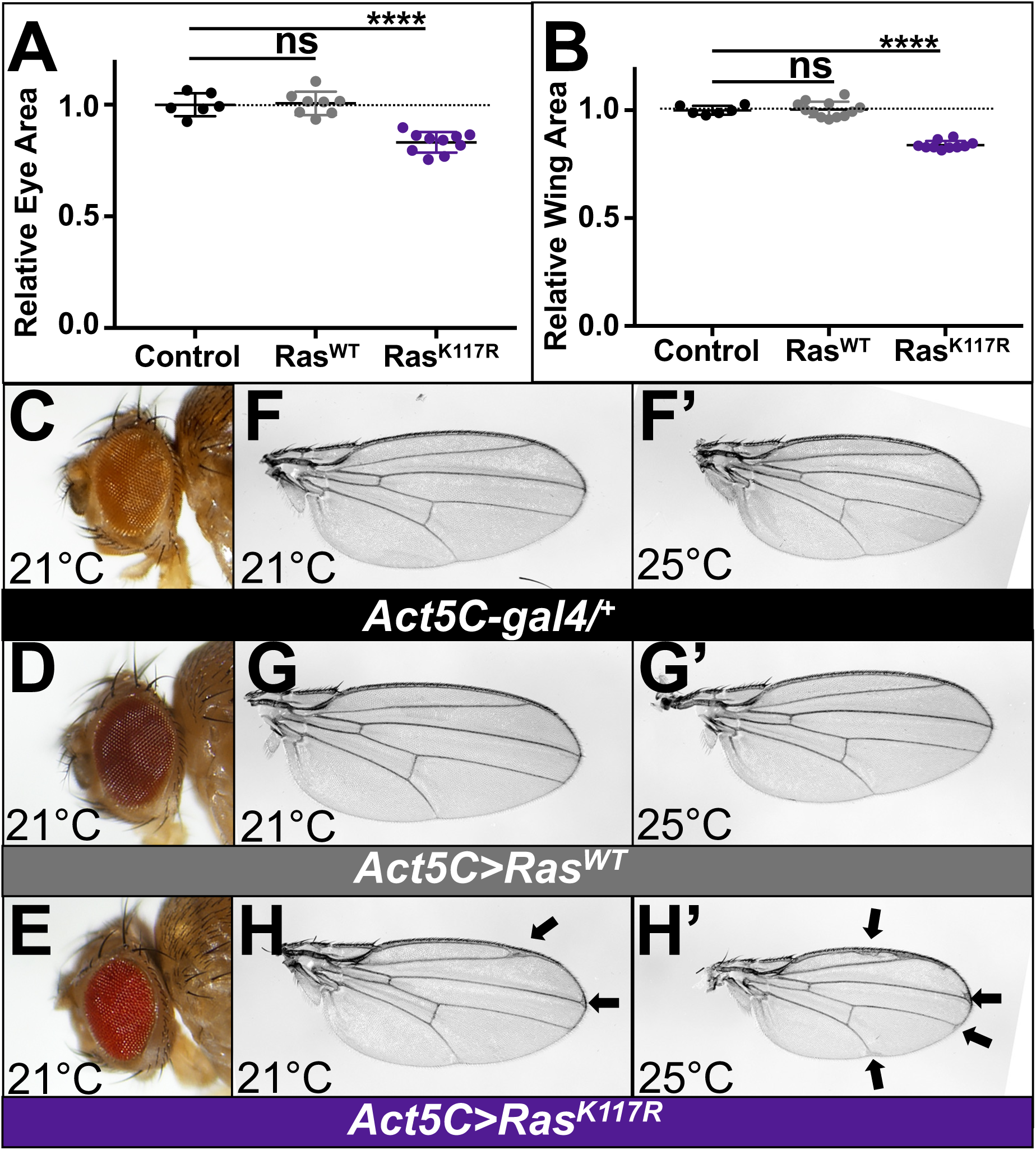
Constitutive expression of Ras^K117R^ reduced organ size and caused rough eye and ectopic wing vein phenotypes. (A) Scatter plot showing the relative eye size of control *Act5C-gal4/+*, *Act5C>Ras^WT^*, and *Act5C>Ras^K117R^*eyes. Constitutively expressing Ras^WT^ (lane 2) did not alter relative eye area compared to *Act5C-gal4/+*controls (lane 1), but constitutively expressing Ras^K117R^ (lane 3) statistically significantly reduced eye size. Sample eyes from this graph are shown in panels C, D, and E. (B) Scatter plot showing the relative wing size of control *Act5C-gal4/+*, *Act5C>Ras^WT^*, and *Act5C>Ras^K117R^*wings. Constitutively expressing Ras^WT^ (lane 2) did not alter relative wing area compared to *Act5C-gal4/+* controls (lane 1), but constitutively expressing Ras^K117R^ (lane 3) statistically significantly reduced wing size. Sample wings from this graph are shown in panels F-H’. Dotted lines overlaid on graphs in in A-B indicate the mean for the control. ns indicates not significant, **** indicates statistically significant by ANOVA; for p values, see Supplemental File S1. (C) Control *Act5C-gal4/+* eye. (D) *Act5C>Ras^WT^*eye. Constitutive Ras^WT^ expression did not cause any obvious eye phenotypes. (E) *Act5C>Ras^K117R^* eye. Constitutive Ras^K117R^ expression causes eyes that were noticeably smaller and rougher than control eyes. Flies in A-E were reared at 21°C. (F-F’) Control *Act5C-gal4/+* wing at (F) 21°C or (F’) 25°C. (G-G’) *Act5C>Ras^WT^* wings at (G) 21°C or (G’)25°C. Constitutive Ras^WT^ expression caused no obvious wing phenotypes. (H-H’) *Act5C>Ras^K117R^* wings at (H) 21°C or (H’) 25°C. Ras^K117R^ expression at (H) 21°C caused phenotypes evident where L2 and L3 meet the wing margin (arrows). These phenotypes worsened at (H’) 25°C and also included ectopic veins anterior to L2 and where L4 and L5 meet the wing margin (arrows). Eyes and wings in A-H’ are from female flies. Female images are shown in panels C-H’ and analyzed in panels A-B.

### Constitutive Ras^K117R^ expression causes eye and wing phenotypes consistent with Ras gain-of-function phenotypes

In addition to size phenotypes, Ras^K117R^-expressing flies showed gross morphological changes in the eye and wing compared to Ras^WT^-expressing or *Act5C-gal4/+* controls (Fig. 2). *Act5C>Ras^K117R^* eyes were obviously rough (Fig. 2E) compared to *Act5C-gal4/+* controls (Fig. 2C) and *Act5C>Ras^WT^* eyes (Fig. 2D). The eye roughness resembled that seen for driving Ras gain-of-function in the eye seen previously for driving a G12V mutationally activated form of Ras in the pattern of the *sevenless* gene or with *GMR-gal4* [Fortini et al., 1992; Karim et al., 1996; Yan et al., 2010; Washington et al., 2020]. *Act5C>Ras^K117R^*wings had ectopic and thickened veins most obvious near the region anterior to L2 and vein deltas or other disrupted pattern where L2, L3, L4, and L5 meet the wing margin (Fig. 2H-2H’). These wing vein phenotypes were not seen upon driving constitutive expression of Ras^WT^ (Fig. 2G-2G’) or in control *Act5C-gal4/+* flies (Fig. 2F-2F’). These phenotypes resembled those seen when driving Ras gain-of-function mutations using *ms1096-gal4* seen previously [Washington et al., 2020; Karunaraj et al., 2024].

### Trametinib and Rigosertib increased survival of K117R expressing flies but did not suppress the reduced organism and organ size phenotypes

To further evaluate this model, we repeated these studies in the presence of Ras pathway inhibitors. Trametinib, a direct MEK inhibitor [Abe et al., 2011; Gilmartin et al. 2011] that therefore inhibits signaling through the Raf-ERK effector branch, statistically significantly increased survival of *Act5C>Ras^K117R^*flies to adulthood compared to a DMSO solvent control (Fig. 3A). This is consistent with lethality resulting, at least in part, from increased signaling through the Raf-ERK branch of Ras signaling. Ras pathway inhibitor Rigosertib [Athuluri-Divakar et al., 2016; Baker et all 2020] also statistically significantly increased survival of *Act5C>Ras^K117R^* flies to adulthood compared to DMSO controls (Fig.3A). Rigosertib has been reported to affect the Raf-ERK branch although some reports suggest this may be indirect [Wang et al., 2021; Ritt et al., 2016], so this would be consistent with the results for Trametinib. Rigosertib has also consistently been described to have potent activity towards Ras effector PI3K [Prasad et al., 2016; Ma et al., 2012; Chapman et al., 2012]. Therefore, this would also be consistent with lethality in part resulting from increased signaling through the PI3K branch.

**Figure 3:**
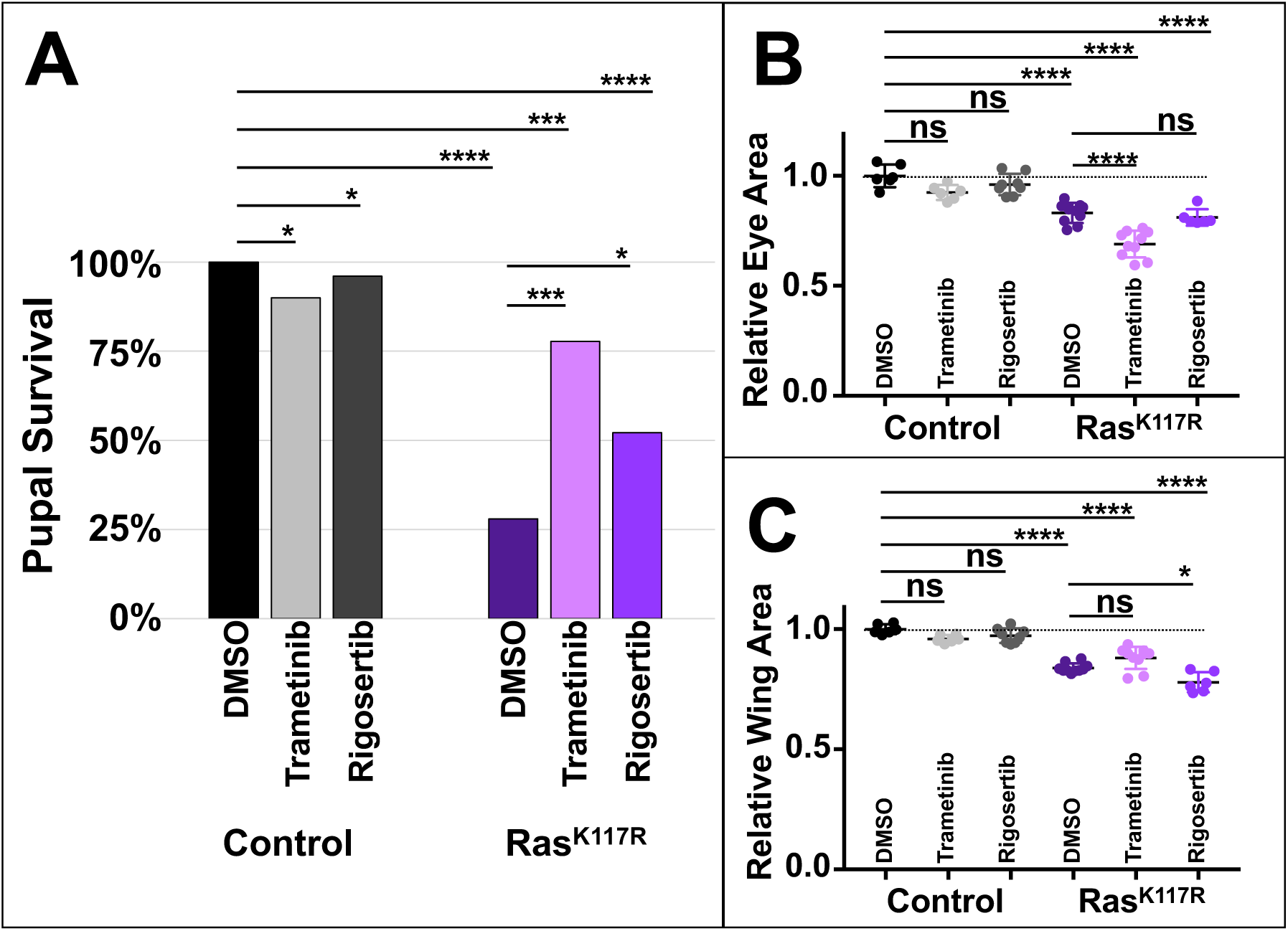
Trametinib and Rigosertib suppressed the lethality of constitutive Ras^K117R^ expression but did not suppress the reduced organ size phenotypes. (A) Bar graph summarizing pupal survival of control *Act5C-gal4/+* flies (lanes 1-3) or Ras^K117R^ flies (lanes 4-6). Treatment with 1 μM Trametinib (lane 2) or μM 10 Rigosertib (lane 3) did not statistically significantly alter survival of control flies compared to a DMSO control (lane 1). Treatment with 1 μM Trametinib (lane 5) or μM 10 Rigosertib (lane 6) statistically significantly increased the survival of constitutive Ras^K117R^ expressing pupae compared to a DMSO control (lane 4), although the survival was still statistically significantly different from *Act5C-gal4/+* controls (lane 1). (B) Scatter plot showing relative eye area for *Ac5Ct-gal4/+* control eyes (lanes 1-3) and Ras^K117R^ eyes (lanes 4-6) treated with Trametinib and Rigosertib. Treatment with 1 μM Trametinib (lane 2) or μM 10 Rigosertib (lane 3) did not affect relative eye size of *Act5C-gal4/+* control flies compared to treatment with a DMSO control (lane 1). Trametinib did not suppress but in contrast statistically significantly enhanced (lane 5) the reduced eye size phenotype of constitutive Ras^K117R^ expression (lane 4). Rigosertib did not affect the reduced eye size phenotype (lane 6). (C) Treatment with 1 μM Trametinib (lane 2) or μM 10 Rigosertib (lane 3) did not affect relative wing size of *Act5C-gal4/+* control flies compared to treatment with DMSO (lane 1). Trametinib did not affect the reduced wing size phenotype (lane 5). Rigosertib did not suppress but in some trials statistically significantly enhanced (lane 6) the reduced wing size phenotype of constitutive Ras^K117R^ expression (lane 4). Experiments in A-C were conducted at 21°C; the baseline eye and wing data graphed in panels B-C were from the same trial also shown in Figures 2. Females were analyzed in panels B-C. Dotted lines overlaid on graphs in in B-C indicate the mean for the control. For graphs in A-C, ns indicates not significant, **** indicates statistically significant by Chi square (A) or by ANOVA (B-C); for p values, see Supplemental File S1.

Despite the striking suppression of lethality, neither Trametinib nor Rigosertib suppressed the reduced size phenotypes of *Act5C>Ras^K117R^* flies (quantified for eye size in Fig. 3B and for wing size in Fig. 3C; images shown in Fig. 4). This would be consistent with the small stature phenotypes resulting from Raf-independent and PI3K-independent branches of Ras signaling. Other *Drosophila* RASopathy models have linked some such phenotypes to other Ras effectors including to downstream regulation of cAMP [Zhong, 1995; Guo et al., 1997; Tong et al., 2002; Walker et al., 2006]. Future work should explore a role for cAMP signaling and other Ras effectors in the small size phenotypes.

**Figure 4:**
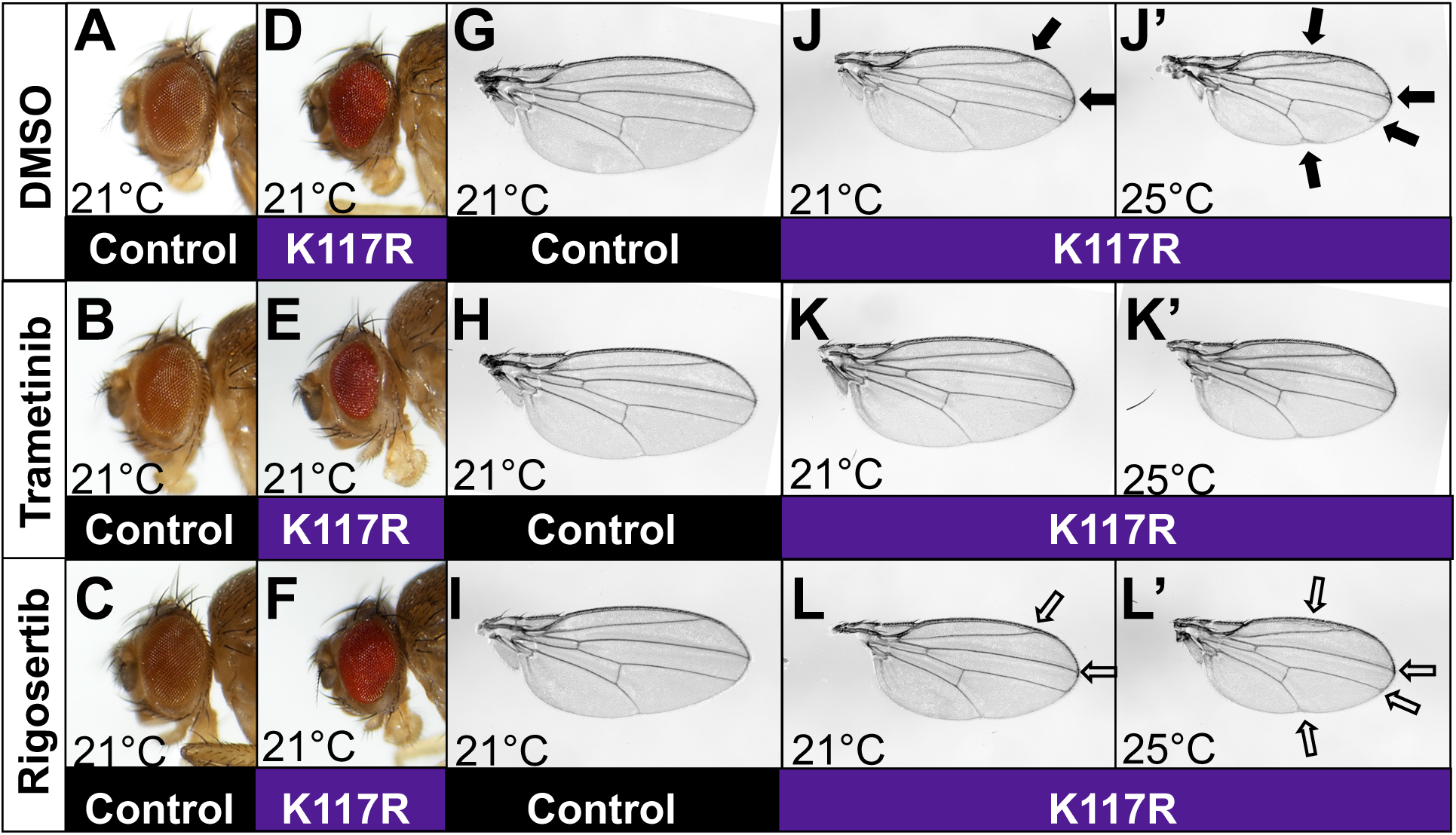
Trametinib and Rigosertib showed different activity for Ras^K117R^ eye and wing phenotypes. (A) Control *Act5C-gal4/+* eye from fly reared on DMSO-food. (B) Control Act5C-gal4/+ eye from fly reared on food containing 1 μM Trametinib. (C) Control *Act5C-gal4/+* eye from fly reared on food containing 10 μM Rigosertib. (D) *Act5C>Ras^K117R^*eye from fly reared on DMSO-containing food. As in Figure 2, the eyes were noticeably smaller and rougher than control eyes. (E) *Act5C>Ras^K117R^* eye from fly reared on food containing 1 μM Trametinib. Trametinib-treated *Act5C>Ras^K117R^* flies had eyes even smaller and rougher eyes than DMSO-fed *Act5C>Ras^K117R^* flies. (F) *Act5C>Ras^K117R^* eye from fly reared on food containing 10 μM Rigosertib. Rigosertib treatment did not noticeably alter the *Act5C>Ras^K117R^* eye phenotype. Flies in A-F were reared at 21°C. (G) Control *Act5C-gal4/+* wing from fly reared on DMSO-food. (H) Control *Act5C-gal4/+* wing from fly reared on food containing 1 μM Trametinib. (I) Control *Act5C-gal4/+* wing from fly reared on food containing 10 μM Rigosertib. (J-J’) *Act5C>Ras^K117R^* wings from flies reared on DMSO-containing food at 21°C or 25°C. As in Figure 2, Ras^K117R^ expression at 21°C caused phenotypes evident where L2 and L3 meet the wing margin (arrows). These phenotypes worsened at 25°C and also included ectopic veins anterior to L2 and where L4 and L5 meet the wing margin (arrows). (K-K’) *Act5C>Ras^K117R^* wings from flies reared on 1 μM Trametinib-containing food at 21°C or 25°C. The wing vein phenotypes were strikingly suppressed. (L-L’) Act5C>Ras^K117R^ wings from flies reared on 10 μM Rigosertib-containing food at 21°C or 25°C. There was some subtle suppression of the wing vein phenotypes (open arrows) as scored by independent lab members, but the phenotypes were still substantial. Female images of eyes and wings are shown in A-L’.

Surprisingly Trametinib reproducibly statistically significantly enhanced the reduced eye size and eye roughness phenotypes of *Act5C>Ras^K117R^*flies (eye size quantified in Fig. 3B; eye morphology shown in Fig. 4E) compared to DMSO-fed *Act5C>Ras^K117R^* controls (Fig. 3B, Fig. 4D). Rigosertib did not affect the reduced eye size phenotypes (size quantified in Fig. 3B; eyes shown in Fig. 4F). Trametinib had no effect on the reduced wing size phenotype of *Act5C>Ras^K117R^*flies (Fig. 3C). In contrast, in some trials, Rigosertib enhanced the reduced wing size of *Act5C>Ras^K117R^*flies (Fig. 3C).

The enhancement of reduced eye size by Trametinib and reduced wing size by Rigosertib is perplexing, as these phenotypes are consistent with Ras effector branches for which Trametinib and Rigosertib have activity. We speculate that the efficacy of these drugs may have been sufficient to disable negative feedback mechanisms which in turn resulted in enhancement of the phenotype. Alternatively, the action of these drugs may have resulted in other signaling outputs that affect size in these contexts for which the mechanism is not yet clear.

### Trametinib suppressed the wing vein phenotypes of Ras^K117R^ expressing flies

Trametinib visibly suppressed the wing vein phenotypes (Fig. 4K-4K’) compared to flies fed a DMSO control (Fig. 4J-4J’). This is consistent with the *Act5C>Ras^K117R^* wing vein phenotypes resulting from increased signaling through the RAF-ERK effector branch. This was an expected result; increased signaling through the ERK branch is associated with wing vein phenotypes.

In contrast, although independent observers scored subtle suppression of wing vein phenotypes by Rigosertib (Fig. 4L-4L’ compared to baseline phenotypes shown in Fig. 4J-4J’), Rigosertib did not cause such striking suppression of wing vein phenotypes as was seen with Trametinib (Fig. 4K-4K’ compared to Fig. 4J-4J’). This result is consistent with reports that Rigosertib primarily has effects on the PI3K branch of Ras signaling [Prasad et al., 2016; Ma et al., 2012; Chapman et al., 2012] with more indirect effects on the ERK branch [Wang et al., 2021; Ritt et al., 2016]. P3K signaling downstream of Ras has not been associated with these wing vein phenotypes.

Together, these data build on previous work supporting the fly system as an excellent model for studying the mechanisms underlying Costello Syndrome phenotypes and for evaluating therapeutic candidates. More specifically, these data support that flies can be used to model Costello Syndrome and colorectal cancers associated with rare K117R mutations.

Curiously, K117 is not only a site of activating mutations in Costello Syndrome and colorectal cancer, but it is also the site of activating ubiquitination [Dard et al., 2022; Kerr et al., 2006; Haigis, 2017]. Whereas K117R appears to confer gain-of-function as a solitary mutation, in the context of activating G12V mutations, we showed previously that adding a second site K117R mutation (to generate Ras^G12V,K117R^) caused much milder phenotypes than Ras^G12V^ alone [Singh et al., 2023]. Although this seems like a paradox, this was presumably due to lack of activating ubiquitination or acetylation events. It will be important for future work to explore further how K117 influences overall Ras activation levels.

## MATERIALS AND METHODS

### Rigor and Reproducibility

The reported work represents reproducible experiments that reflect a minimum of three well-controlled, independent trials. For phenotypes that are subjective (not quantifiable), independent lab members score progeny blinded to genotype to avoid bias. To avoid replicating unintended observer bias, for all experiments, at least one trial was conducted by a different lab member.

### Pupal and adult lethality/survival experiments

For pupal lethality experiments, pupal cases of the indicated genotypes were scored as dead (in which dead pupae remained in the pupal cases) or empty (from which surviving flies had eclosed) and counted as in our previous study [Singh et al., 2023] by 34 days (18°C) 26 days (21°C) or 18 days (25°C). Dead pupal cases are easily distinguished from empty pupal cases or from developing pupae that are still alive and have not yet eclosed. We also scored adult flies from crosses in which we recorded the number of male and female flies of each genotype as indicated by the presence or absence of balancer markers.

### *UAS Ras^K117R^* transgenic line

*UAS Ras^K117R^ was* cloned into pUAST-attB using the EcoRI and NotI restriction sites. FLAG and His6 sequences MDYKDDDDKRGSHHHHHHALE were added immediately after the EcoRI site (corresponding to N-terminal nucleotide sequence GAATTCATGGATTACAAGGATGACGACGATAAGAGAGGATCGCATCACCATCACCATCACGCGCTCGAG, EcoRI site underlined) as we did previously with UAS Ras^G12V^ and various Y4 mutants [Washington et al., 2020; Karunaraj et al., 2024], and for Ras constructs in the G12V background with second site mutations at K117 and K147 [Singh et al. 2023]. An additional stop codon and a NotI site were added immediately after the original stop codon (corresponding to the sequence TAATAAGCGGCCGC, original stop codon underlined). The plasmids were sent to BestGene for injection. The resulting line was balanced with second chromosome balancers and allowed to homozygose. Homozygous lines were maintained as true-breeding homozygous stocks. BestGene used techniques for site-specific integration at attp40, and we sequenced genomic DNA to confirm the desired K117R insert sequence. However, the eye color of the transgenic line used in this work is darker than the other lines made similarly, so there may be a second insertion closely linked to attp40 also in this line. This new transgene is listed in Table 1.

**Table 1:** Table of reagents used with corresponding identifiers.

| ***Drosophila* Strains** |  |  |
| --- | --- | --- |
| **Strain** | **Source** | **Identifier** |
| *w<sup>1118</sup>* | The fly community and Bloomington *Drosophila* Stock Center (BDSC) | BL-3605, BL-5905 and others<br>RRID:BDSC_3605, RRID:BDSC_5905 |
| *Act5C-gal4* | BDSC | Can be obtained from BDSC, BL-3954<br>RRID:BDSC_3954 |
| *UAS Ras<sup>WT</sup>* | Washington et al., 2020 |  |
| *UAS Ras<sup>K117R</sup>* | This study |  |
| **Software** |  |  |
| **Program** | **Source** |  |
| Image J | https://imagej.net/ij/ |  |
| GraphPad Prism | https://www.graphpad.com/scientific-software/prism/ |  |
| Microsoft Excel | https://www.microsoft.com/Microsoft/Excel/ |  |

### *Drosophila* husbandry and general experiment information

Crosses were set up at the indicated temperatures using standard *Drosophila* medium as in our previous works [Yan et al., 2009; Washington et al., 2020; Reimels et al., 2024]. For each experimental trial, crosses used food prepared in the same batch and were incubated in close proximity to ensure the same environment to rule out unintended environmental variables or variations in food sources. We cannot rule out subtle environmental differences or differences in batches of food used in different trials that may have contributed to variability between trials. *Act5c-gal4* driver was obtained from the Bloomington Stock center (details, Table 1). The *UAS Ras^K117R^*transgene was generated for this study as described, and the *UAS Ras^WT^* transgene was first reported in our previous work [Washington et al., 2020]. We scored experiments for both males and females, but given the increased lethality of *Act5C>Ras^K117R^* males, not every trial had sufficient males for analysis, so the adult eye and wing analysis was performed on females.

### Image analysis and processing

Adults, adult wings, and adult were photographed using a Nikon DS-Fi3 microscope camera and saved as TIFF files. Raw images were converted to grayscale in Adobe Photoshop. Brightness and contrast of wing images were adjusted in Adobe Photoshop for clarity; all adjustments were applied across the entire images. Genotypes are summarized and identifiers are annotated in Table 1.

### Drug treatment

Trametinib (GSK1120212) was purchased from Fisher Scientific and added to the fly food for a final concentration of 1μM as has been used in other *Drosophila* studies [Levine et al., 2016; Bangi et al., 2019]. Rigosertib was kindly provided by Drs. Stacey Baker and Prem Reddy and was added to the fly food for a final concentration of 10μM. To add the drugs to food, we used two different methods: (1) standard fly food was melted, allowed to cool to 50°C upon which the drug or DMSO solvent was added. (2) Carolina formula 4-24 blue flakes were reconstituted with liquid containing either DMSO solvent or drug according to 0.75 grams of food in 3 mls of liquid. Both food sources resulted in the same reproducible phenomena. Data shown in Figures 3-4 reflect one of the three trials that used method (1) for standard *Drosophila* media. Control phenotypes shown in Figures 1-2 are from flies reared in DMSO control food.

### Statistical analysis

The relative length of the adult notum and abdomen and the relative area of the adult eye and wing area were measured using ImageJ software in pixels, then normalized using Excel, and finally graphed and analyzed using t tests in GraphPad Prism (Fig. 1F-1G) or one way ANOVA in GraphPad Prism (Fig. 2A-2B, Fig. 3B-3C) as appropriate. Pupal or adult counts for lethality experiments were graphed in Excel and analyzed based on categories (dead/eclosed or Males/females) in GraphPad Prism using contingency analysis for categorical scoring in Chi square tests or Fisher’s exact tests (Fig. 1A-1A’, Fig. 3A). P values from Chi square tests are shown in Figures, P values from Chi square tests and Fisher’s exact test are listed in Supplemental File S1.

### Genotypes of flies in images and graphs

*Act5C-gal4/+* (Fig. 1B, left fly in Fig. 1E, Fig. 2C, Fig. 2F-2F’, Fig. 4A-4C, Fig. 4G-4I; left-most lane in graphs in Fig. 1A, Fig. 1A’, Fig. 1F, Fig. 1G, Fig. 2A, Fig. 2B, left-most three lanes of Fig. 3A, Fig. 3B, Fig. 3C)

*Act5C>UAS Ras^WT^* (Fig. 1C, Fig. 2D, Fig. 2G-2G’, middle lanes in Fig. 1A, Fig. 1A’, 2A, Fig. 2B)

*Act5C>UAS Ras^K117R^* (Fig. 1D, right fly in Fig. 1E, Fig. 2E, Fig. 2H-2H’, Fig. 4D-4F, Fig. 4J-L’; right lanes in Fig. 1A, Fig. 1A’, Fig. 1F-1G, Fig. 2A-2B, right-most three lanes Fig. 3A, Fig. 3B, Fig. 3C)

## DATA AVAILABILITY STATEMENT

*Drosophila* strains used in this work (listed in Table 1) have been published previously [Washington et al., 2020], are available from public stock centers, or are available upon request. Raw data, normalized data, and p values for graphs in Figs. 1-3, are listed in Supplemental File S1. The authors affirm that all data necessary for interpreting the data and drawing conclusions are present within the article text, the figures, table, and Supplemental File S1.

## ACKNOWLEDGMENTS

We thank M Mlodzik, U Weber, TK Das, J Chipuk, P Rangan, ZQ Pan, and the New York Fly community. We thank K. Kalafsky, K. Braden, E. Loizides, T. Zuluaga, T. Hyunh, R. Chernet, C. Washington, C. Ye for assistance. We thank the Bloomington Drosophila Stock Center (NIH P40OD018537) for providing fly stocks and Flybase (NIH 5U41HG000739) for access to sequence information.

